# *AtGPP2* encodes a 3-deoxy-*manno-*octulosonate-8-phosphatase required for the synthesis of KDO in rhamnogalacturonan II

**DOI:** 10.64898/2026.05.02.722439

**Authors:** Tomomi Hara, Yinglin Wang, Masaru Kobayashi, Toru Matoh

## Abstract

3-Deoxy-D-*manno*-oct-2-ulosonic acid (KDO) is an essential component of rhamnogalacturonan II (RG-II), a complex pectic polysaccharide required for plant growth and development. While most steps of the KDO biosynthetic pathway have been characterized in plants, KDO-8-phosphatase (KDO8Pase), the phosphatase responsible for converting KDO 8-phosphate (KDO8P) to KDO, remained unidentified. To identify this missing component, we performed gene co-expression analysis and identified At5g57440 (GPP2) as the primary candidate in Arabidopsis (*Arabidopsis thaliana* L.). Recombinant GPP2 protein exhibited KDO8P-specific phosphohydrolase activity in vitro. A GFP-tagged GPP2 protein was predominantly localized to mitochondria, consistent with the compartmentation of the subsequent step in KDO biosynthesis. Null mutants of *GPP2* exhibited significant growth retardation under boron-limited conditions, in which expression of *GPP2* and other KDO biosynthetic genes was up-regulated. The growth retardation was also observed in liquid culture in normal media, a condition that induces rapid growth and thus likely increases metabolic demand for KDO. Despite this growth defect, the KDO content per unit cell wall in *gpp2* remained equivalent to that in wild-type plants. These results are consistent with the identification of GPP2 as the elusive plant KDO8Pase and suggest a model where KDO availability becomes the rate-limiting factor for cell wall production. Our findings complete the plant KDO biosynthetic pathway and provide new insights into the physiological significance of RG-II in cell wall biosynthesis.

**Significance statement:** This study identifies the previously unknown plant KDO-8-phosphatase, thereby completing the biosynthetic pathway for KDO in rhamnogalacturonan II. Our findings demonstrate that KDO synthesis is up-regulated under boron deficiency, and its supply becomes a rate-limiting factor for cell wall formation.

## Introduction

3-Deoxy-D-*manno*-oct-2-ulosonic acid (KDO) is an eight-carbon sugar acid found in Gram-negative bacteria and vascular plants. In plants, KDO occurs as a residue of rhamnogalacturonan II (RG-II), a domain of pectin in the primary cell wall (York *et al*., 1985). RG-II forms a dimer, being cross-linked with boric acid (Kobayashi *et al*., 1996; O’Neill *et al*., 1996), thereby binding pectin macromolecules to form a hydrophilic gel that surrounds cellulose microfibrils. The gel is considered crucial for maintaining the structural integrity of the cell wall (Ryden *et al*., 2003) and is indispensable for normal growth and development (O’Neill *et al*., 2004).

Although RG-II is a relatively small polysaccharide consisting of about 30 sugar residues, it contains 12 different constituent sugars, including rare sugars such as aceric acid, apiose, 3-deoxy-D-*lyxo*-heptulosaric acid (DHA), KDO, 2-*O*-methyl-L-fucose or 2-*O*-methyl-D-xylose (O’Neill *et al*., 2004). Previous studies have shown that disruption of the synthesis of RG-II-specific sugars leads to severe growth defects, indicating that each of these unique glycosyl residues plays an essential role in the proper function of RG-II (Hays *et al*., 2025). KDO has also been demonstrated to be indispensable for plant growth, development, and fertility (Kobayashi *et al*., 2011; Delmas *et al*., 2008; Séveno *et al*., 2010; Zhang *et al*., 2024). However, the precise contribution of KDO to RG-II function remains largely unknown.

Analysis of mutants defective in KDO biosynthesis would be useful for clarifying the physiological role of KDO in RG-II, and such mutants can be obtained by perturbing the expression of genes for the KDO biosynthetic pathway. In Gram-negative bacteria, KDO occurs as a component of extracellular lipopolysaccharides, and its biosynthetic pathway has been extensively characterized (Cipolla et al., 2010; Raetz and Whitfield, 2002). The KDO biosynthesis in plants is hypothesized to follow a pathway similar to that of bacteria (Smyth and Marchant, 2013): D-arabinose 5-phosphate (Ara5P), generated from D-ribulose 5-phosphate by D-arabinose-5-phosphate isomerase (API), is condensed with phosphoenolpyruvate (PEP) by KDO-8-phosphate synthase (Kdos) to produce KDO 8-phosphate (KDO8P). KDO8P is hydrolyzed by a specific phosphatase to yield KDO, which is subsequently activated to cytidine 5’-monophosphate KDO (CMP-KDO) by cytidine 5’-triphosphate:KDO cytidylyltransferase (CMP-KDO synthetase; CKS). The formed CMP-KDO serves as the substrate for KDO transferase. Several genes involved in this pathway have already been identified (Fig. 1). The Arabidopsis (*Arabidopsis thaliana* L.) gene *SETH3* (*At3g54690*) (Lalanne *et al*., 2004) encodes a protein similar to *Escherichia coli* API (Meredith and Woodard, 2003). *At1g79500* and *At1g16340* encode Kdos-like proteins, whereas *At1g53000* encodes a CKS-like protein, and their enzymatic activities have been experimentally confirmed (Matsuura *et al*., 2003; Misaki *et al*., 2009; Kobayashi *et al*., 2011). *MGP2/RCKT1* (*At1g08660*) was originally identified as a gene whose mutation causes defects in pollen function (Deng *et al*., 2010). This gene, as well as its homolog *SIA2* (*At3g48820*), was predicted to encode an enzyme transferring KDO and/or DHA based on sequence similarity to animal sialyltransferases (Deng *et al*., 2010; Dumont *et al*., 2014). A recent study has provided evidence supporting that *MGP2/RCKT1* functions as a KDO transferase acting on RG-II (Zhang *et al*., 2024). Meanwhile, the specific phosphatase responsible for converting KDO8P to KDO has remained a “missing link” in plants. In *E. coli*, the existence of such phosphatase [KDO-8-phosphatase (KDO8Pase); EC 3.1.3.45] was first suggested by Ghalambor *et al*. (1966), subsequently purified by Ray and Benedict (1980), and assigned to the *KdsC* (formerly known as *YrbI*) gene by Wu and Woodard (2003). Given the conservation of other steps between bacterial and plant KDO biosynthesis (Smyth and Marchant, 2013), it is reasonable to assume that plant genomes also encode phosphatase(s) specific for KDO8P. Nevertheless, simple homology searches using the bacterial KdsC sequence as a query fail to identify any highly similar plant proteins, suggesting that plants may possess a functionally equivalent but structurally divergent enzyme that catalyzes this reaction.

**Figure 1.**
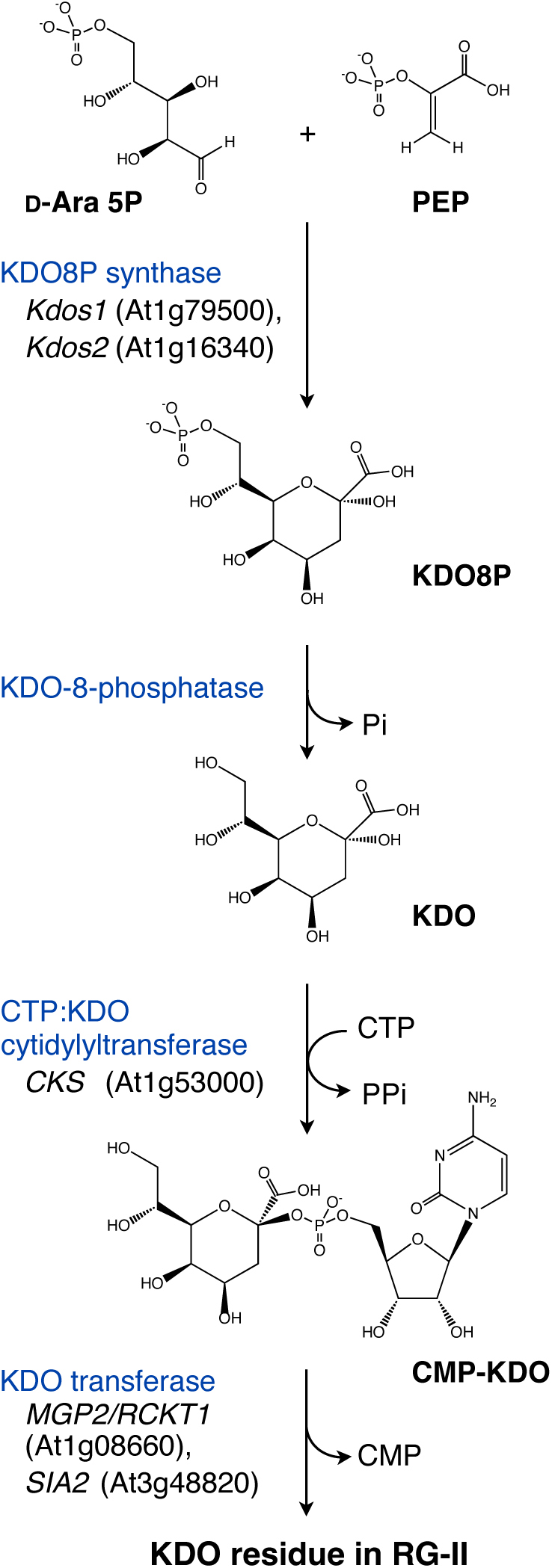
Predicted biosynthetic pathway of RG-II KDO residue in plants. Enzymes and corresponding genes identified in *Arabidopsis thaliana* are indicated.

To identify the gene encoding plant KDO8Pase, we employed gene co-expression analysis in Arabidopsis and identified *At5g57440* (*GPP2*) as the primary candidate. Here, we demonstrate that GPP2 is a mitochondria-localized phosphatase with specificity for KDO8P. Phenotypic evaluation of *gpp2* null mutants reveals that their growth is impaired under the conditions of increased KDO demand, and the KDO availability becomes a rate-limiting factor for cell wall production under such conditions. Our findings complete the plant KDO biosynthetic pathway and provide new insights into the physiological significance of RG-II in cell wall biosynthesis.

## Results

### Identification of candidate gene by gene co-expression analysis

In our attempt to identify the protein catalyzing the hydrolysis of KDO8P in Arabidopsis, we first performed a BLAST search using the bacterial KdsC sequence as the query but could not find any protein with sufficiently high similarity. We, therefore, employed a gene co-expression analysis using the ATTED-II database (http://atted.jp) (Obayashi *et al*., 2022) to identify the genes showing an expression pattern similar to those of genes involved in KDO biosynthesis in Arabidopsis. Table I shows the top 10 genes showing highly correlated patterns of expression with *Kdos1* (*At1g79500*), which encodes an isoform of two KDO8P synthases in *A. thaliana* (Matsuura *et al*., 2003). *Kdos2*, the gene encoding another isoform of KDO8P synthase in Arabidopsis, showed the highest co-expression score (Table 1). The list also contains the gene encoding a sialyltransferase-like protein, MGP2/RCKT1, a Golgi-localized KDO transferase (Deng *et al*., 2010; Zhang *et al*., 2024) (Table 1). The occurrence of these known or putative KDO-related genes in the list suggested the usefulness of this approach as a way to search genes encoding KDO8Pase. We then focused on At5g57440, ranked second on the list, as the primary candidate gene encoding KDO8Pase in Arabidopsis. The gene encodes a hydrolase-like protein that has been previously reported as a homolog of the yeast glycerol-3-phosphatase and designated *GPP2* (Caparrós-Martín *et al*., 2007).

**Table 1.** The top 10 co-expressed genes with *Kdos1* as searched on ATTED-II database.

| Rank | Gene | Predicted function | Coexpression Z-score |
| --- | --- | --- | --- |
| 1 | At1g16340 | KDO8P synthase 2 | 8.66 |
| 2 | At5g57440 | Haloacid dehalogenase-like hydrolase superfamily protein | 8.17 |
| 3 | At3g20790 | NAD(P)-binding Rossmann-fold superfamily protein | 6.00 |
| 4 | At3g09800 | SNARE-like superfamily protein | 5.91 |
| 5 | At5g64860 | Disproportionating enzyme | 5.83 |
| 6 | At3g07200 | RING/U-box superfamily protein | 5.75 |
| 7 | At5g49555 | FAD/NAD(P)-binding oxidoreductase family protein | 5.69 |
| 8 | At4g17190 | Farnesyl diphosphate synthase 2 | 5.61 |
| 9 | At3g12290 | Amino acid dehydrogenase family protein | 5.50 |
| 10 | At1g08660 | MGP2 (sialyltransferase-like protein) | 5.28 |

The AlphaFold2-predicted structure of GPP2 (AF-Q8VZP1 in AlphaFold Protein Structure Database) exhibits a typical HAD enzyme architecture, consisting of a Rossmann fold–like core domain and a cap domain (Burroughs *et al*., 2006) (Fig. 2A). At the amino acid sequence level, GPP2 showed only limited similarity to KdsC, the bacterial KDO8Pase (Fig. 2C). However, the structure of the core domain, which is presumed to be responsible for catalytic activity, was highly similar to that of KdsC (Fig. 2B). For this comparison, we used the KdsC from *Moraxella catarrhalis* (McKdsC), whose crystal structure has been determined in a ligand-bound form (PDB ID: 4UMF) (Dhindwal *et al*., 2015), so that structural information on the active site could be obtained. Molecular docking simulations indicated that GPP2 can accommodate KDO8P within the putative substrate-binding pocket with a configuration in which KDO8P is predicted to form hydrogen bonds and electrostatic interactions with several nearby residues (Fig. 3). In addition, consistent with the ubiquitous presence of KDO as a constituent residue of RG-II in vascular plants, sequences highly similar to GPP2 were identified across a wide range of plant species, including dicots, monocots, and gymnosperms (Fig. 4). Since these features are consistent with those expected for a plant KDO8Pase, we further characterized this gene and its product in the following analyses.

**Figure 2.**
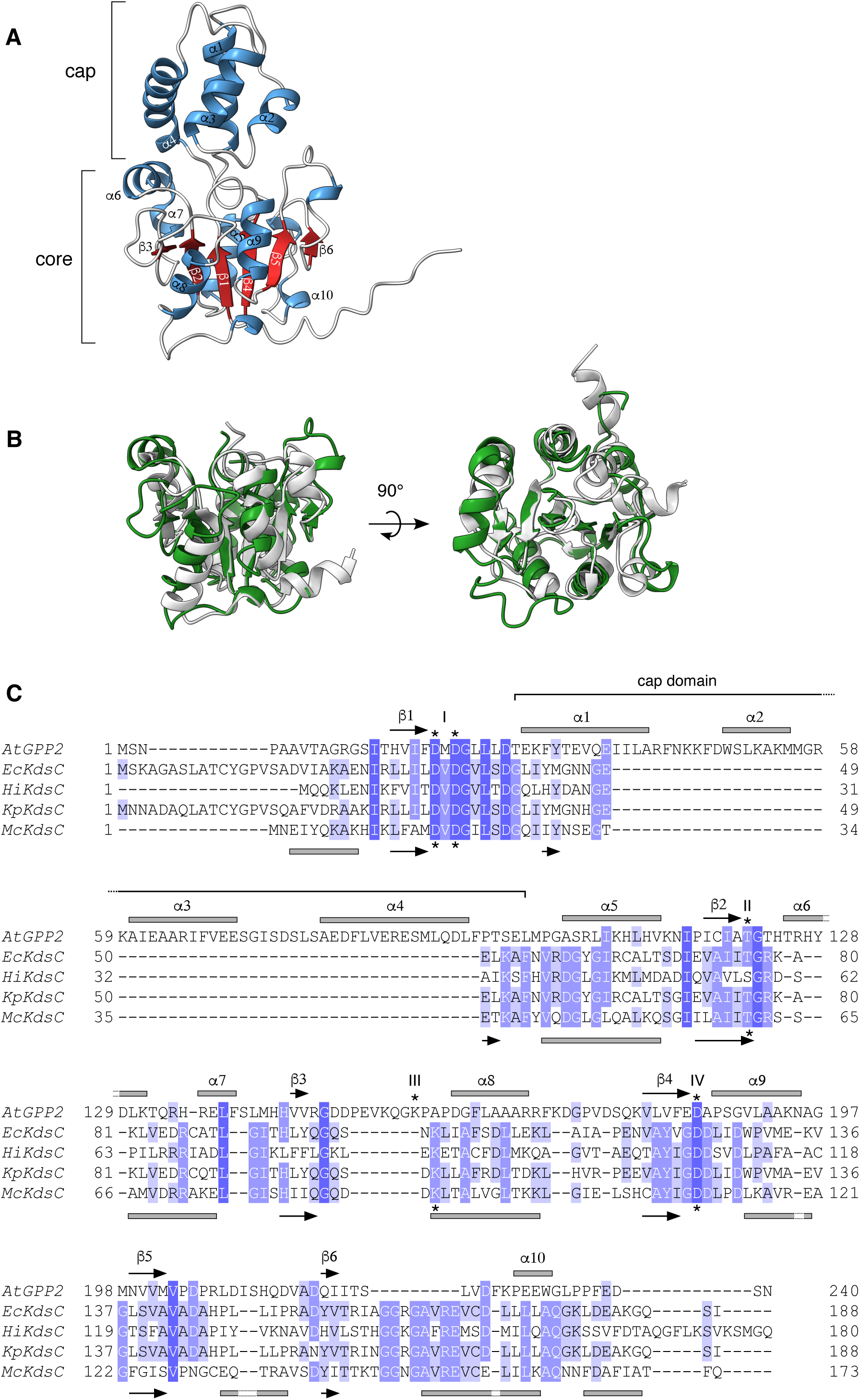
Structure of GPP2 protein. (A) The AlphaFold2-predicted structure of GPP2 (AF-Q8VZP1). Helices and β-strands are shown in blue and red, respectively. Labels on the secondary structures correspond to those in Fig. 3. (B) Superposition of the core domain of GPP2 (green) and KdsC, the bacterial KDO8Pase from *Moraxella catarrhalis* (gray). The core domain of GPP2 was aligned with the core region of McKdsC (PDB ID 4UMF), excluding a large β-hairpin protruding from the core (Q26-A38). (C) Sequence comparison of GPP2 with bacterial KDO8Pases. The Arabidopsis GPP2 sequence was aligned with bacterial KDO8Pases, including KdsC proteins from *Escherichia coli* (PDB ID: 3HYC), *Haemophilus influenzae* (1J8D), *Klebsiella pneumoniae* (7T35), and *Moraxella catarrhalis* (4UMF). Secondary structure elements are indicated above and below the alignment for AtGPP2 and McKdsC, respectively, with ɑ-helices shown as boxes and β-strands as arrows. Residues marked with asterisks correspond to the conserved HAD motifs (I–IV).

**Figure 3.**
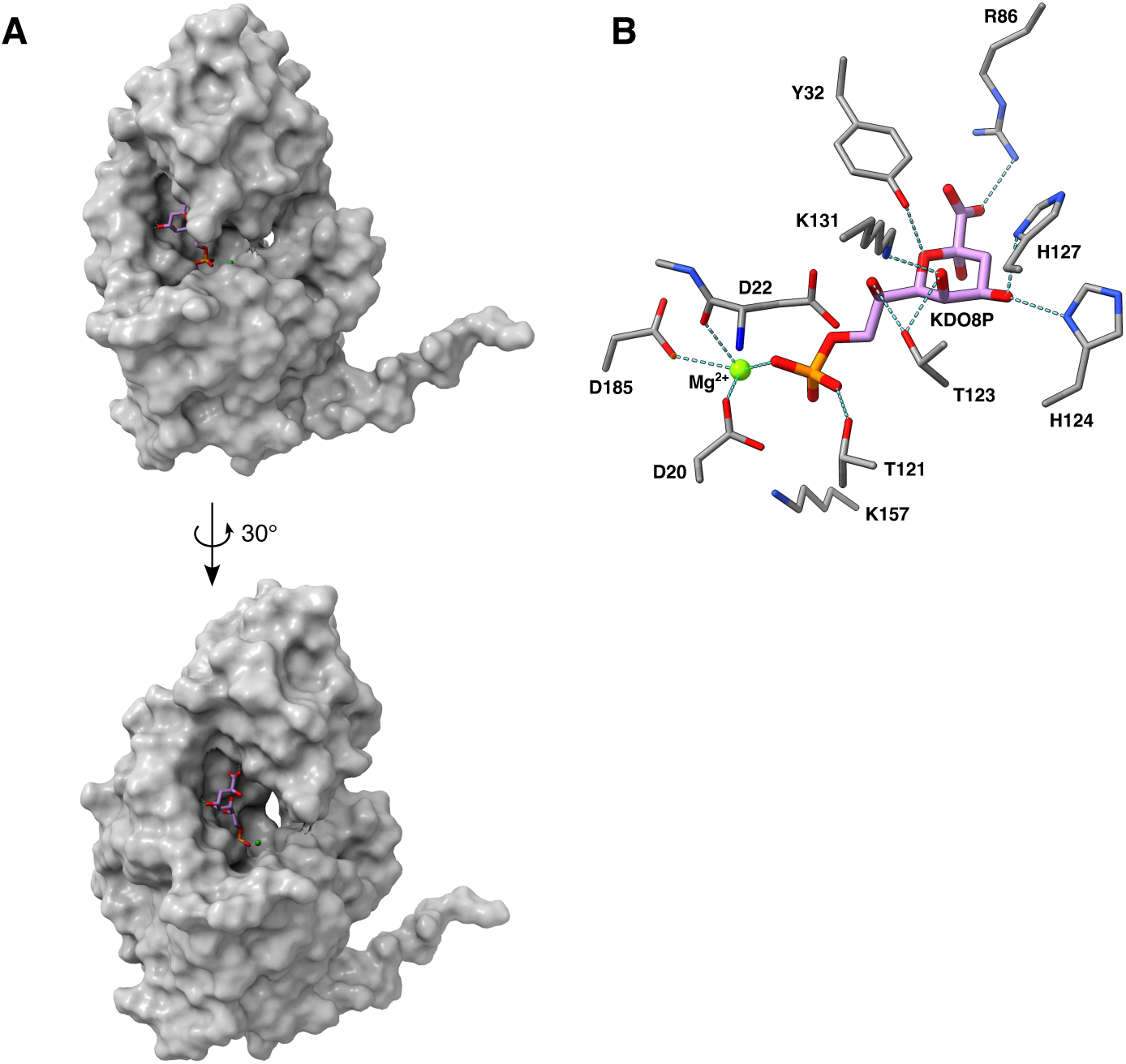
Active site model of GPP2 with KDO8P. (A) Surface representation of GPP2 with KDO8P shown as sticks in a modeled binding mode based on docking simulation and manual refinement. The view corresponds to that in Fig. 2A, with a rotation to improve visualization of the putative active site. (B) Close-up view of the putative active site. Residues potentially involved in substrate recognition and catalysis are indicated. Putative interactions, including coordination with Mg^2+^, electrostatic interactions, and hydrogen bonds, are shown as dashed lines.

**Figure 4.**
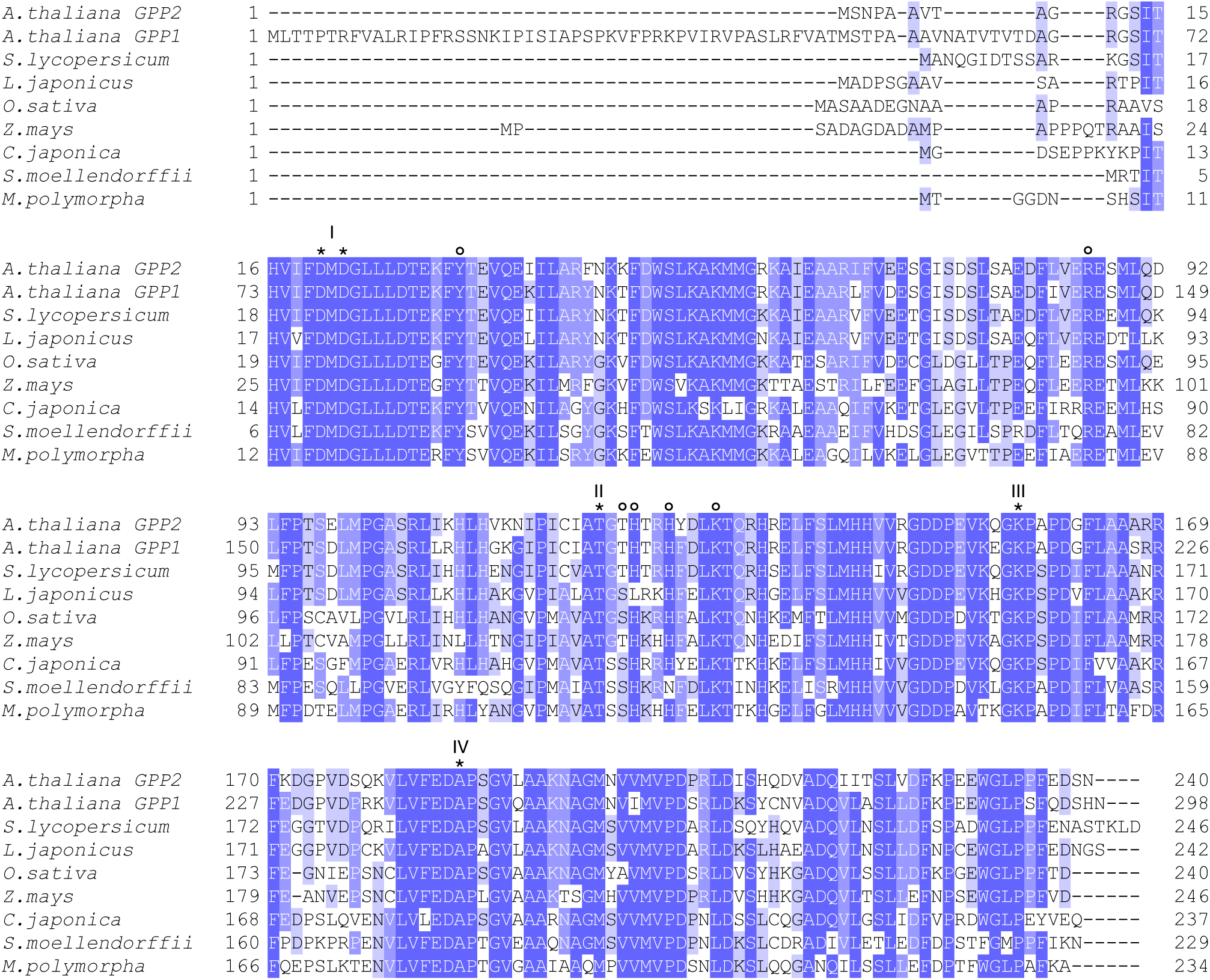
Alignment of plant GPP homologs. Multiple sequence alignment of GPP proteins from representative plant species, including Arabidopsis GPP1 and GPP2, *Solanum lycopersicum* XP_004248690, *Lotus japonicus* XP_057421206, *Oryza sativa* XP_015648459, *Zea mays* NP_001353093, *Selaginella moellendorffii* XP_002970198, *Marchantia polymorpha* Mp3g13310, *Cryptomeria japonica* IABU01028871. Residues marked with asterisks correspond to the conserved HAD motifs (I–IV), while residues indicated by open circles represent those predicted to interact with KDO8P, as shown in Fig. 3B.

### Enzymatic characterization of recombinant GPP2 protein

We first examined whether the GPP2 protein could hydrolyze KDO8P. A recombinant protein with the full-length GPP2 protein fused to the C-terminus of glutathione-*S*-transferase (GST) was expressed in *E. coli* and purified using a glutathione affinity resin (Fig. 5A). When the fusion protein was mixed with KDO8P, a time-dependent release of inorganic phosphate (Pi) was observed (Fig. 5B). On the other hand, heat-inactivated protein or GST alone did not cause the release of Pi (data not presented), indicating that the GPP2 portion of the fusion protein hydrolyzed KDO8P. As GPP2 was previously proposed to be a glycerol-3-phosphatase (Caparrós-Martín *et al*., 2007), we examined the activity of the GST–GPP2 fusion protein on glycerol 3-phosphate as well. However, in our hands, the fusion protein did not show any detectable activity toward glycerol 3-phosphate (Fig. 5B). No activity was observed for fructose 6-phosphate, D-arabinose 5-phosphate, PEP, or *p*-nitrophenyl phosphate (data not presented). These results together suggest that the GPP2 protein is a KDO8P-specific phosphohydrolase. As expected for a putative HAD enzyme, the KDO8Pase activity of GST–GPP2 required divalent cations, including Mg^2+^ and Mn^2+^ (Fig. 5C). In the presence of Mg^2+^, the activity was highest around pH 8 (Fig. 5D).

**Figure 5.**
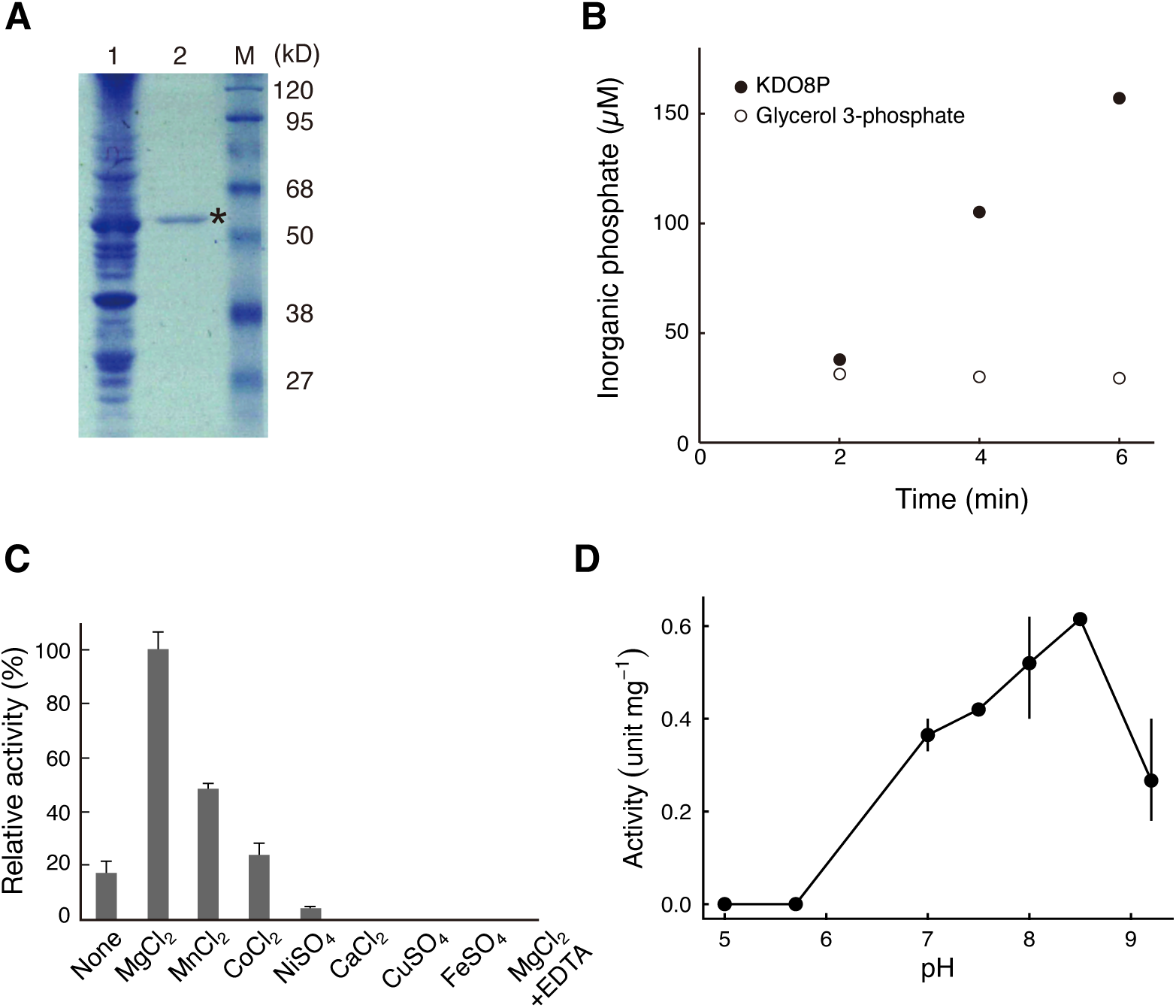
Enzymatic characterization of recombinant GPP2 protein. (A) Purification of recombinant GPP2 protein. Crude lysate of *E. coli* cells expressing the recombinant GST–GPP2 protein (lane 1) and the fraction purified by glutathione affinity chromatography (lane 2) were electrophoresed on a 10% SDS-polyacrylamide gel. Lane M: molecular weight markers. The asterisk indicates the band corresponding to GPP2. (B) Time course of Pi production from KDO8P and glycerol 3-phosphate. A representative result from three independent experiments is shown. Reactions were performed in 25 mM Tris–HCl (pH 8.0) containing 5 mM MgCl_2_ and 0.3 mM KDO8P or 1 mM glycerol 3-phosphate. (C) Metal requirement for the activity. Reactions were performed in 25 mM Tris–HCl (pH 8.0) containing 0.3 mM KDO8P and the indicated additives. Activities are shown as relative to the activity observed in the presence of Mg^2+^. Data points and vertical lines indicate the mean and range of the data from two replicates, respectively. (D) pH dependence of the activity. Data points and vertical lines indicate the mean and range of the data from two or three replicates, respectively.

The Arabidopsis genome encodes another protein, GPP1 (At4g25840; Caparrós-Martín et al., 2007), which shows high sequence similarity to GPP2. The deduced amino acid sequence of GPP1 contains an approximately 50-residue N-terminal extension that is absent from GPP2 (Fig. 4) and is predicted to function as a targeting signal sequence (Caparrós-Martín et al., 2007). Excluding this region, GPP1 shares 83% sequence identity with GPP2 at the protein level. We therefore examined the KDO8Pase activity of GPP1 as well. A truncated form of GPP1 lacking the first 50 amino acid residues was expressed in *E. coli* as a GST-fusion protein. The GST–GPP1 fusion protein catalyzed the release of Pi from KDO8P, but not from glycerol 3-phosphate, as observed for GST–GPP2 (Fig. 5A). Kinetic analysis revealed that GST–GPP2 and GST–GPP1 exhibited similar *V*max values, whereas GST–GPP2 showed a slightly lower *K*m value for KDO8P than GST–GPP1 (Table 2).

**Table 2.** Kinetic parameters of recombinant GPP1 and GPP2 proteins.

| Protein | $K_m$ (mM) | $V_{max}$ (U mg<sup>-1</sup>) |
| --- | --- | --- |
| GST-GPP1 | 1.10 ± 0.003 | 4.6 ± 2.7 |
| GST-GPP2 | 0.219 ± 0.134 | 3.8 ± 0.35 |

Values represent means ± SD from three independent experiments.

### Subcellular localization of GPP proteins

To investigate the subcellular localization of GPP1 and GPP2 proteins, green-fluorescent protein (GFP) was fused to the C-terminus of full-length proteins, and the fusion proteins were transiently expressed in Arabidopsis seedlings. Since the expression levels were either too high or too low when using the cauliflower mosaic virus 35S (CaMV35S) RNA promoter or their own promoters, respectively, we employed *ubiquitin 10* (*UBQ10*) promoter (Grefen *et al*., 2010) in this study. The GPP2–GFP fusion protein exhibited a punctate fluorescence pattern (Fig. 6A). When the seedlings expressing the fusion protein were fed with MitoTracker dye to stain mitochondria, co-localization of MitoTracker and GFP fluorescence was observed (Fig. 6A), suggesting localization of GPP2 protein in mitochondria. In contrast, fluorescence from the GPP1–GFP fusion protein was mostly co-localized with chlorophyll autofluorescence (Fig. 6B). The result suggests that GPP1, unlike GPP2, localizes to plastids.

**Figure 6.**
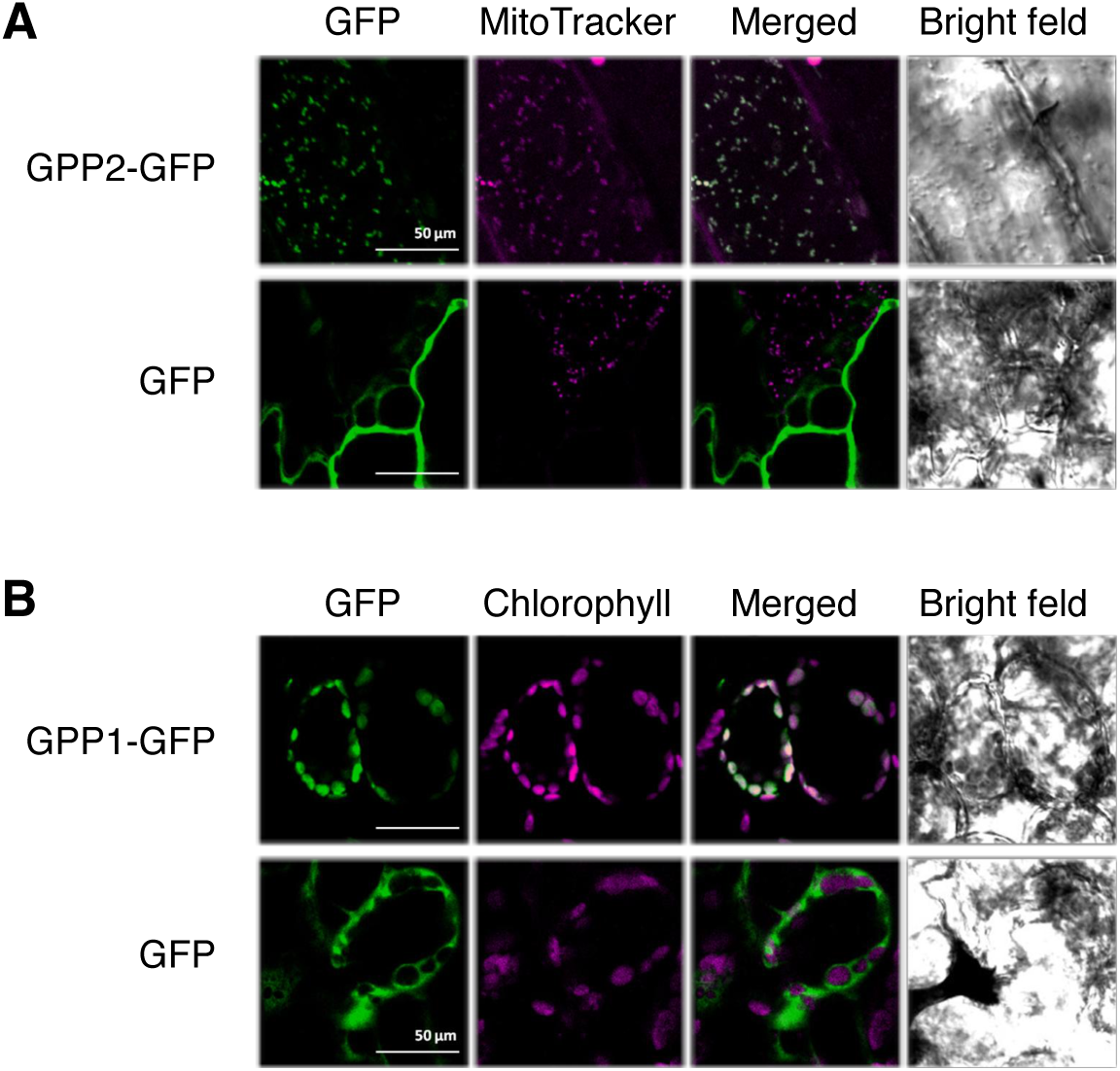
Subcellular localization of GPP proteins. Confocal laser scanning microscopy images of Arabidopsis cells expressing GFP, GPP1–GFP, or GPP2–GFP. GFP fluorescence, MitoTracker staining, chlorophyll autofluorescence, and merged images are shown. (A) GPP2–GFP or GFP alone. (B) GPP1–GFP or GFP alone.

### Knockout mutants of GPP2 show boron-dependent phenotype

To evaluate the physiological significance of GPP2 proteins, we characterized plants lacking GPP2. Out of three independent T-DNA insertion lines obtained from the Arabidopsis Biological Resource Center (ABRC), SALK_062406 and SALK_017815 were found to be *GPP2* null mutants (Fig. 7). Hereafter, the lines are referred to as *gpp2-1* (SALK_062406) and *gpp2-2* (SALK_017815).

**Figure 7.**
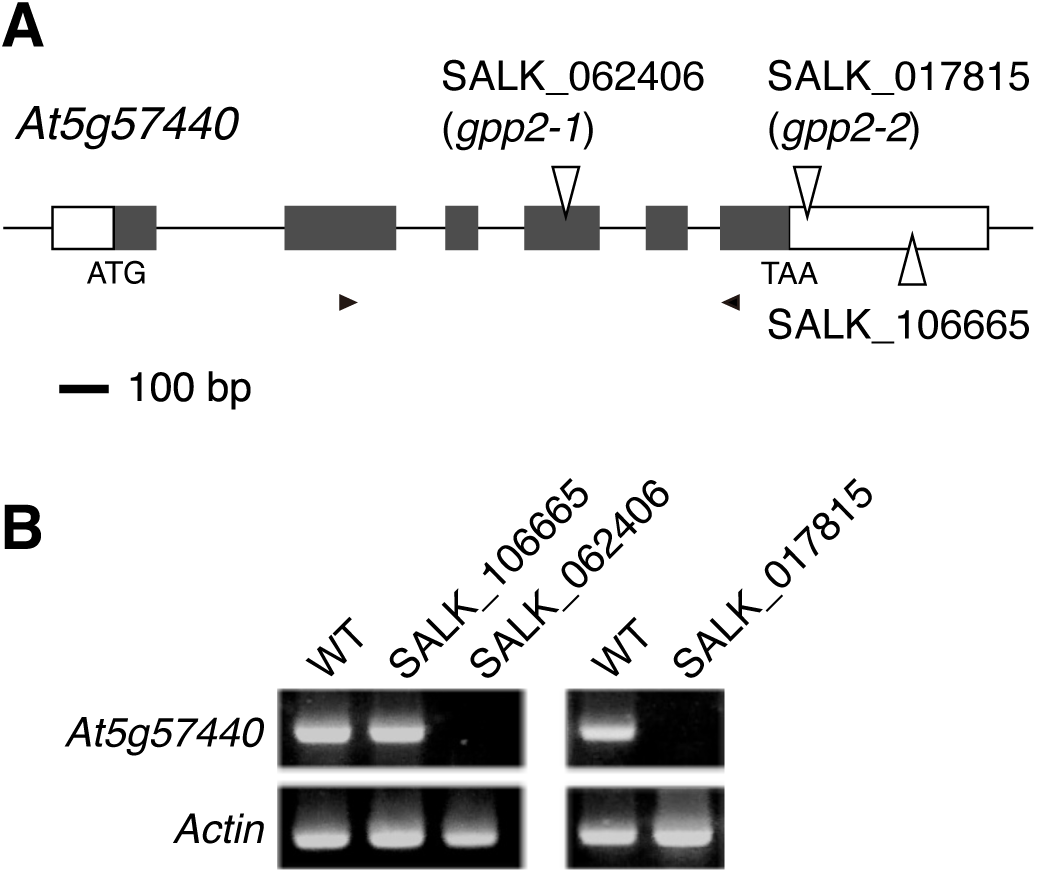
Schematic representation of the *GPP2* gene (At5g57440) and T-DNA insertion sites in the allele. Boxes represent exons and solid lines represent introns. Gray boxes represent protein-coding sequence, and arrowheads indicate the annealing site of primers. (A) Reverse transcription–PCR analysis of *GPP2* transcript abundance in T-DNA insertion lines. Total RNA was extracted from young leaves of either WT or T-DNA mutant plants, reverse transcribed, and PCR amplified. *Actin 2* (At3g18780) was included in the analysis to calibrate cDNA concentrations.

Both the *gpp2-1* and *gpp2-2* mutants showed no discernible phenotype when grown on a half-strength Murashige-Skoog (MS) solid medium supplemented with vitamins and sucrose. Meanwhile, their primary roots became shorter than wild-type (WT) plants under a boron (B)-limited condition (0.3 µM) (Fig. 8A). Other stress treatments, including salinity (30 mM NaCl) and low pH (pH 4), did not cause significant differences in growth between the mutants and WT (Supplementary Fig. S1). When plants were cultured in vitro in half-strength liquid MS media supplemented with vitamins and sucrose, the growth of mutants was reduced compared with WT, even under normal B supply (Fig. 8B). Meanwhile, the contents of KDO and B per unit cell wall weight were not different between mutant and WT (Table 3).

**Figure 8.**
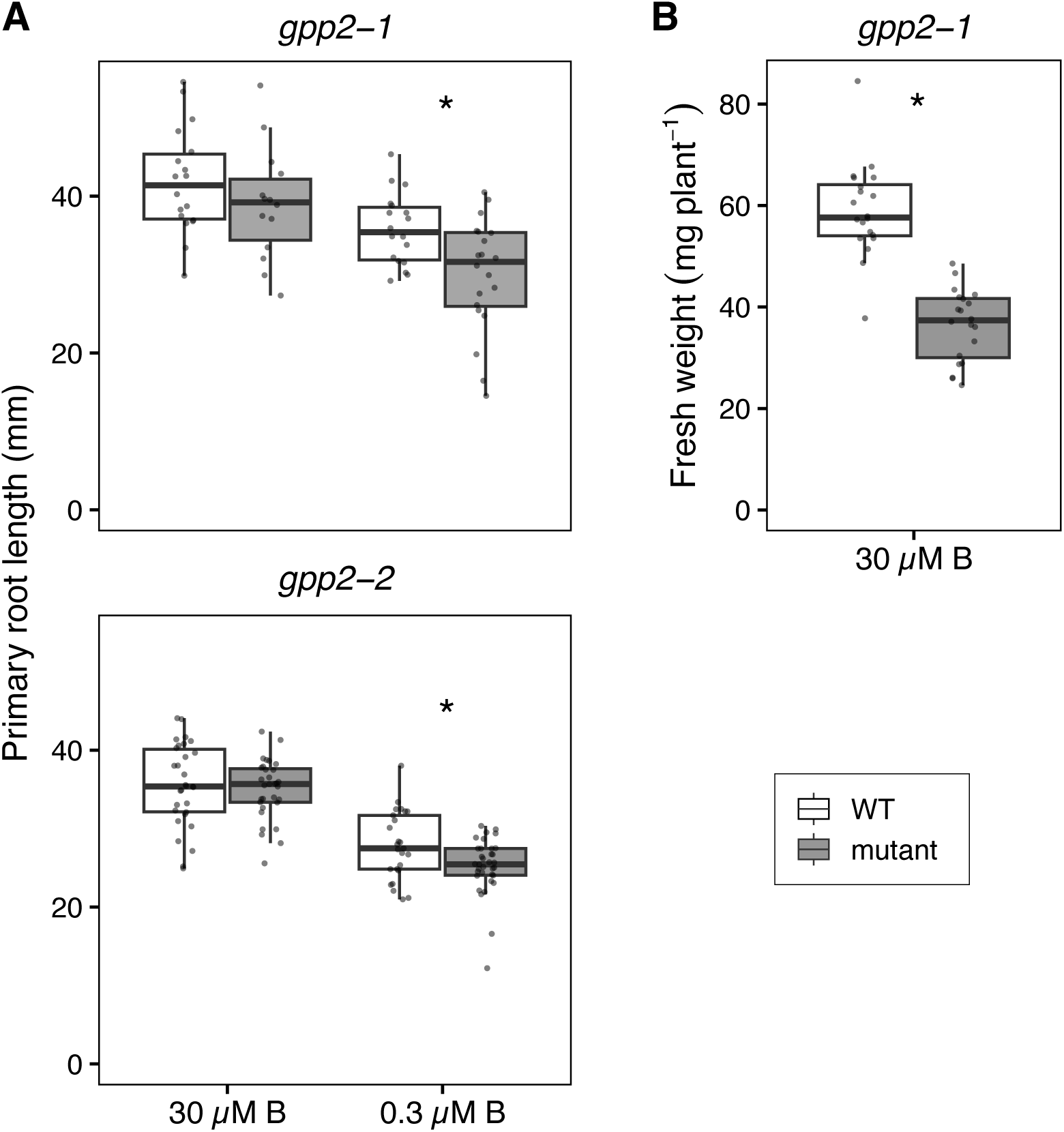
Growth of *gpp2* mutant plants. (A) Primary root length of wild-type (WT) and *gpp2-1* (upper panel) or WT and *gpp2-2* (lower panel) seedlings grown on normal media (half-strength MS solid media containing 30 µM B) or low-B media (0.3 µM B) for 7 days. (B) Fresh weight of WT and *gpp2-1* seedlings grown in a half-strength MS liquid medium under normal B supply (30 µM B) for 11 days. Individual data points are shown on the boxplots. Asterisks indicate significant differences between genotypes (P < 0.05; Student’s *t*-test; n = 20–33 for A and n = 20 for B).

**Table 3.**
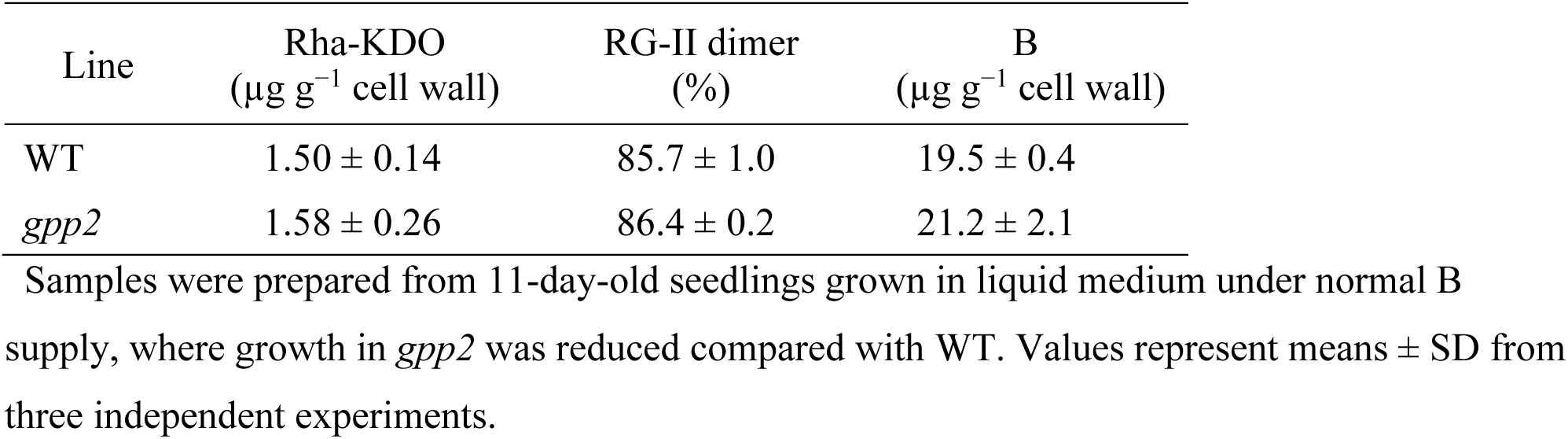
Comparison of the cell walls of WT and *gpp2* mutant plants.

| Line | Rha-KDO<br>($\mu\text{g g}^{-1}$ cell wall) | RG-II dimer<br>(%) | B<br>($\mu\text{g g}^{-1}$ cell wall) |
| --- | --- | --- | --- |
| WT | $1.50 \pm 0.14$ | $85.7 \pm 1.0$ | $19.5 \pm 0.4$ |
| *gpp2* | $1.58 \pm 0.26$ | $86.4 \pm 0.2$ | $21.2 \pm 2.1$ |

Samples were prepared from 11-day-old seedlings grown in liquid medium under normal B supply, where growth in *gpp2* was reduced compared with WT. Values represent means ± SD from three independent experiments.

To explore the causal relationship between the *GPP2* mutation and the growth phenotype, we examined the expression of *GPP2* under low-B conditions. As shown in Fig. 9, more *GPP2* transcripts were detected under the low-B condition than in the normal condition. We also examined the expression of known or putative KDO synthesis-related genes and found that the expressions of *Kdos* (Matsuura *et al*., 2003) and *CKS* (Kobayashi *et al*., 2011) were also up-regulated under the low-B condition (Fig. 9). *GPP1* expression was not induced under this condition (Fig. 9). These results suggest that the synthesis of KDO residues in RG-II is up-regulated under B-limited conditions.

**Figure 9.**
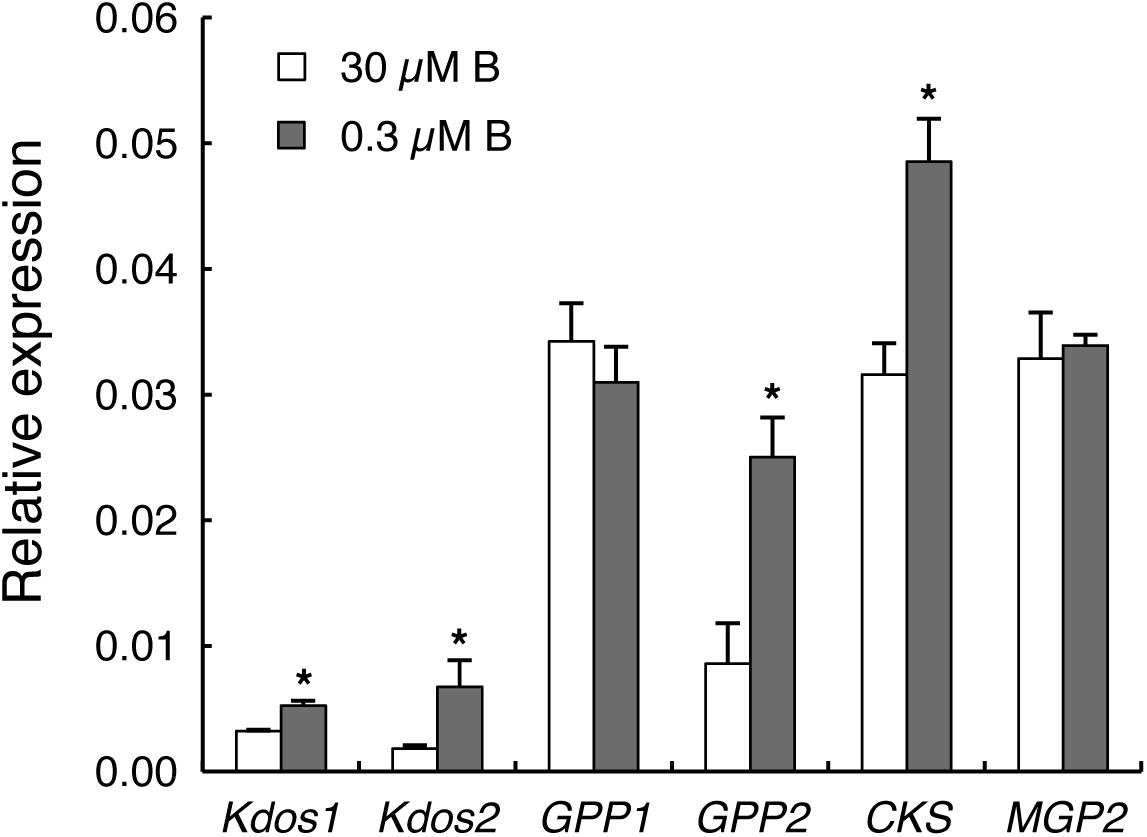
Expression of *GPP2* and other KDO-related genes under low-B conditions. Transcript levels were quantified by RT-qPCR in 5-d-old seedlings grown on normal (30 µM B) or low-B (0.3 µM B) medium. Relative expression levels were normalized to *actin 2*. Bars and vertical lines indicate the means and SD, respectively. Asterisks indicate significant differences between the conditions (*p* < 0.05; Student’s *t*-test; n = 3).

## Discussion

In this study, we aimed to identify the gene encoding KDO8Pase in plants and characterized the product of *GPP2*, a gene showing highly correlated expression with known KDO-related genes in Arabidopsis (Table 1). Consistent with the ubiquitous presence of RG-II in vascular plants, sequences closely similar to GPP2 were identified across a wide range of vascular plant species (Fig. 4). Interestingly, closely related sequences were also found in a bryophyte, *Marchantia polymorpha* (Fig. 4), in which the presence of RG-II has not been confirmed (Hays et al. 2025). Liverworts are known to produce secondary metabolites containing KDO derivatives (Katayama *et al*., 2012), raising the possibility that the GPP2-like protein may be involved in the biosynthesis of KDO derivatives in such compounds.

The GPP2 recombinant protein exhibited phosphohydrolase activity toward KDO8P (Fig. 5B), but not toward other organic phosphates examined. A previous study has reported GPP2 as a glycerol-3-phosphatase (Caparrós-Martín *et al*., 2007). However, in our hands, the recombinant GPP2 protein did not show any detectable phosphohydrolase activity on glycerol 3-phosphate (Fig. 5B). At present, we cannot exclude the possibility that this discrepancy might be attributed to differences in the experimental systems; we expressed the GPP2 protein in *E. coli* as a GST fusion protein, while the previous study used maltose-binding protein as the fusion partner (Caparrós-Martín *et al*., 2007). Nonetheless, KDO8P seems to be a better substrate for this enzyme than glycerol 3-phosphate; the apparent *K*_m_ values were 0.22 mM for KDO8P (Table 2) and 3.5 mM for glycerol 3-phosphate (Caparrós-Martín *et al*., 2007), and the *V*_max_ values were 3.8 U mg^−1^ protein for KDO8P (Table 2) and in the range of tens mU mg^−1^ protein for glycerol 3-phosphate (Caparrós-Martín *et al*., 2007). The observed *K*_m_ value for KDO8P was comparable to that of *E. coli* YrbI (75 µM; Wu and Woodard 2003), whereas the *V*_max_ value was substantially lower than that of YrbI (500 U mg^−1^; Wu and Woodard 2003). For Arabidopsis Kdos, which catalyzes the upstream step in KDO biosynthesis, a specific activity of 0.827 U mg^−1^ has been reported under fixed substrate concentrations (3 mM PEP and D-arabinose 5-phosphate) (Matsuura *et al*., 2003). Although this value does not represent a *V*_max_ and thus does not allow a direct comparison with GPP2, it suggests that the observed activity of recombinant GPP2 (Table 2) is not incompatible with a role in the KDO biosynthetic pathway.

GPP2 protein was predicted as a member of the HAD superfamily. Most HAD proteins are characterized by a bipartite architecture consisting of a catalytic core domain and a cap domain, the latter of which is typically involved in determining substrate specificity (Burroughs *et al*., 2006; Lahiri *et al*., 2006). GPP2 conforms to this canonical structure, and its core domain exhibits a high degree of structural similarity with the catalytic core of bacterial KDO8Pases (Fig. 2B). In contrast, bacterial KDO8Pases lack an intrinsic cap domain within their monomeric units, but an adjacent monomer within the oligomeric complex functions as a “pseudo-cap” in trans (Biswas *et al*., 2009). This structural divergence makes it difficult to assess whether the substrate pocket of GPP2 is structurally equivalent to that of bacterial KDO8Pases. Nevertheless, in silico docking of KDO8P into the AlphaFold2-predicted model of GPP2, guided by the substrate coordinates observed in McKdsC (Dhindwal *et al*., 2015) and related HAD phosphatases, revealed that the putative active site of GPP2 has sufficient spatial volume to accommodate KDO8P (Fig. 3A). In addition, in the modeled configuration, multiple interactions are predicted to be formed between conserved residues of GPP2 (Fig. 4) and KDO8P, including a potential electrostatic interaction between the C1 carboxyl group of KDO8P and R86, or hydrogen bonds between multiple hydroxy groups and ring oxygen and Y32, T123, H124, H127, and K131 (Fig. 3B). These structural insights are consistent with the hypothesis that GPP2 functions not as a generic phosphatase but as a specialized phosphatase that recognizes KDO8P as the substrate.

Regarding the subcellular localization of GPP2, both cytosolic and mitochondrial localization are plausible, considering that the preceding step of KDO biosynthesis catalyzed by Kdos is likely cytosolic (Delmas *et al*., 2008) while the subsequent activation of KDO by CKS occurs in mitochondria (Kobayashi *et al*., 2011). In this study, the GPP2–GFP fusion protein localized to mitochondria (Fig. 6A). The N-terminal region of GPP2 contains a short stretch of approximately 10 amino acids that appears not to be integrated into the core structure in the AlphaFold model (Fig. 2A). According to SUBA5 (https://suba.live) (Hooper *et al*., 2017), GPP2 is predicted to localize to the cytosol; however, predictions from individual algorithms are inconsistent and span multiple compartments, suggesting that a clear targeting signal may be absent. Previous proteomic studies have reported detection of GPP2 in the cytosol (Ito *et al*., 2011; McBride *et al*., 2019), plasma membrane (Miki *et al*., 2019), and mitochondrial proteomes (Heazlewood *et al*., 2004). In addition, immunoblot analysis using an antibody recognizing both GPP1 and GPP2 detected signals in cytosolic and plastid fractions, although the purity of these fractions was not described (Caparrós-Martín *et al*., 2007). Notably, plastid fractions obtained by differential centrifugation are often contaminated with mitochondria, making it difficult to assess plastid localization from these data alone. Taken together, previous evidence regarding the subcellular localization of GPP2 is inconsistent, and further validation is required.

To further validate GPP2 as the Arabidopsis KDO8Pase, we performed a phenotypic analysis of *gpp2* mutant plants. Mutations in known KDO biosynthesis genes impair pollen function; consequently, homozygous mutants are not obtained for the genes, as has been shown with *Kdos* (Delmas *et al*., 2008), *CKS* (Kobayashi *et al*., 2011), and *MGP2/RCKT1* (Deng *et al*., 2010). In contrast, homozygous T-DNA insertion lines for *GPP2* were obtained (Fig. 7), and the null mutant lines exhibited no discernible phenotype when grown on a half MS solid medium (Fig. 8A, with 30 µM B supply). However, their growth was inferior to that of WT when grown under low-B conditions (Fig. 8A, with 0.3 µM B supply) or in a liquid medium (Fig. 8B). Analysis of the cell wall prepared from the liquid-cultured plants (Fig. 8B) revealed that the KDO content per unit cell wall was not reduced in *gpp2* (Table 3). Therefore, the observed growth defect is unlikely to be due to structural defects in RG-II. Instead, it is possible that the formation of normal RG-II containing KDO became a rate-limiting factor for cell wall production, thereby limiting the total amount of biomass synthesized without altering the composition of cell walls. The reason why KDO synthesis was not completely abolished even in the null mutants might be attributed to functional compensation by GPP1. GPP1 possesses an N-terminal extension of approximately 50 amino acids that is predicted to function as a plastid transit peptide. Consistent with this, SUBA5 predicts plastid localization for GPP1. However, Sa *et al*. (2016) reported that a GPP1::GFP fusion protein localized to mitochondria in Arabidopsis. The recombinant GPP1 exhibited KDO8Pase activity (Table 2). Hence, if both GPP1 and GPP2 are localized to mitochondria, GPP1 may partially compensate for the loss of GPP2 function. Although GPP1 expression was not induced under low-B conditions (Fig. 9), its basal activity may still provide sufficient KDO8Pase function under normal conditions but may become insufficient when demand for KDO increases. Future phenotype characterization of *gpp1*/*gpp2* double mutants is expected to provide further insights into their redundancy. Since no T-DNA insertion line for *GPP1* is available from the ABRC, it will be necessary to generate and analyze a deletion or knockdown mutant. Alternatively, the possibility of KDO8P hydrolysis by other phosphatases with low substrate specificity cannot be excluded. Although KDO8P hydrolysis via such pathways might be less efficient than that mediated by a specialized KDO8Pase, it may nevertheless supply a sufficient amount of KDO to support growth under normal conditions (Fig. 8).

The conditional manifestation of the phenotype in *gpp2* might be due to differences in KDO requirements. The primary roots of plate-grown *gpp2* mutants became shorter than those of WT under B-limited conditions (Fig. 8A). The condition also up-regulated the expression of *GPP2* and other KDO-related genes, including *Kdos* and *CKS* (Fig. 9). Previous studies have also reported an increased expression of *Kdos* in *Brassica napus* under B deficiency at mRNA (Pan *et al*., 2012) or protein (Wang *et al*., 2010) level. Since the induced expression of these genes was not observed under low pH or salinity (Supplementary Fig. S1), it is unlikely to be a general stress response. Instead, these results suggest a link between the B-dependent phenotype and the inhibited synthesis of KDO, a component of RG-II. Given that RG-II is the binding site for B, synthesizing more components of RG-II under B-limited conditions may appear paradoxical. One possible explanation is that Arabidopsis plants may upregulate RG-II synthesis to increase the likelihood of encounters between B and RG-II. Boric acid and RG-II can form borate diester spontaneously (Kobayashi et al. 1999; O’Neill et al. 1996), and the monomeric and B-crosslinked dimeric RG-II would be in equilibrium. Thus, supplying more monomeric RG-II may shift the equilibrium toward forming dimeric RG-II, even at the same B concentration. Such a response might occur if Arabidopsis plants sense the B deficiency-induced destabilization of the pectic network rather than the B concentration itself. Whatever the reason, under conditions of increasing demand for KDO, hydrolysis of KDO8P by mechanisms other than GPP2 may be insufficient, leading to the phenotype in *gpp2* (Fig. 8A).

In the case of liquid culture, the growth of *gpp2* was inferior to WT even under normal B supply (Fig. 8B). Under our experimental conditions, Arabidopsis seedlings grew faster in liquid medium than on solid medium. Faster growth requires accelerated cell wall formation, which requires RG-II with the KDO residue as an indispensable component. Therefore, liquid culture is a system that may exaggerate the impact of defects in the KDO synthetic pathway. Taken together, these considerations allow us to interpret the observed phenotype of *gpp2* as being due to suppression of KDO synthesis.

Complementation of the *gpp2* phenotype by an exogenous supply of KDO would provide further evidence for the identity of GPP2 as KDO8Pase. We therefore tried to determine whether supplementing KDO in the plate medium alleviated the low-B-inducible *gpp2* phenotype, as examining liquid culture with the available amount of KDO was difficult. When 0.5 mM KDO was added to the low B-plate medium containing 30 µM B, the primary root length was not different between WT and *gpp2* mutant (Supplementary Fig. S2). However, the commercially available KDO preparation used in the experiment contained B at 27.5 ng mg^−1^ KDO. Adding the KDO preparation at 0.5 mM increased the medium B concentration to 0.6 µM, which falls within the borderline region for whether the low B-inducible *gpp2* phenotype appears. Hence, it is difficult to determine whether the disappearance of the difference between WT and *gpp2* upon the addition of KDO was ascribed to the complementation of *GPP2* mutation by KDO or to an increase in medium B concentration.

In summary, this study has identified GPP2 as the KDO8Pase in Arabidopsis, thereby completing the biosynthetic pathway for the KDO residue in RG-II. Mutant plants of the gene showed a B-dependent phenotype, and its expression was up-regulated under B deficiency. To our knowledge, this is the first report to suggest environmental effects on RG-II biosynthesis. Further study of this gene will contribute to a better understanding of KDO and RG-II biosynthesis and their physiological functions in plants.

## Materials and Methods

### Plant materials

*Arabidopsis thaliana* (L.) Heynh. were grown under a 16 h/8 h light/dark cycle in a growth chamber at 22°C unless otherwise stated. Ecotype Columbia-0 was used as the WT.

### Expression and characterization of recombinant GPP proteins

The full-length GPP2 and the GPP1 lacking the N-terminal 50 amino acid residues were expressed as GST fusion proteins. The primers used in this study are listed in Supplementary Table S1. The *GPP2* cDNA fragment flanked by *Eco*RI and *Xho*I restriction sites was prepared by PCR using *Eco*RI-linked primer 1 and *Xho*I-linked primer 2, then subcloned into the *Eco*RI/*Xho*I site of pGEX-5X-1 protein expression vector (GE Healthcare). A partial cDNA fragment of *GPP1* was PCR-amplified from an Arabidopsis cDNA library using primers 3 and *Not*I-linked primer 4. The amplified fragment, which contained an internal *Sal*I site present at 150 bp downstream to the start codon, was digested with *Sal*I and *Not*I and inserted into the *Sal*I/*Not*I site of pGEX-5X-2 protein expression vector (GE Healthcare). Constructed plasmids were introduced into *E. coli* BL21 for protein expression after verifying the sequence integrity.

Protein expression was induced with 0.5 mM isopropyl-β-D-thiogalactopyranoside (IPTG) at 16°C for 12 h. Bacterial cells were lysed with 1 mg ml^−1^ lysozyme and 0.2% (w/v) Triton X-100 in 10 mM Tris–HCl (pH 8.0), and the recombinant proteins were purified with GSTrap 4B column (GE Healthcare) according to the manufacturer’s instructions. In a typical experiment, the recombinant protein from a 30-ml culture was recovered in a 1-ml column eluate.

KDO8Pase activity of the recombinant proteins was assayed by following the production of Pi at 30°C in a 200-µl reaction mixture containing 25 mM Tris–HCl, pH 8.0, 5 mM MgCl_2_, varying concentrations of KDO8P, and the purified enzyme preparation. For metal requirement assays, MgCl_2_ was omitted or replaced with various divalent metal salts at a final concentration of 5 mM. Reactions were initiated by adding the enzyme. At predetermined time intervals, 50-µl aliquots of the mixture were withdrawn and mixed in microplate wells with 150-µl color reagent (1:6 mixture of 10% (w/v) ascorbic acid and 0.42% (w/v) (NH_4_)_6_Mo_7_O_24_·4H_2_O in 0.5 M H_2_SO_4_) (Ames, 1966). After incubation at 45°C for 20 min, the absorbance at 650 nm was read using iMark plate reader (Bio-Rad). The kinetic parameters (*K*_m_ and *V*_max_) were estimated by nonlinear regression fitting of the Michaelis-Menten equation using the nls function in R (version 4.5.0; R Core Team, 2025).

KDO8P was prepared enzymatically from Ara5P and PEP. A maltose-binding protein-AtKdos fusion protein was expressed and purified as described (Matsuura *et al*., 2003), except that the protein was expressed at 16°C. A reaction mixture containing 50 mM Tris–HCl, pH 8.0, 100 mM KCl, 1 mM DTT, 2 mM PEP, 2 mM Ara5P, and 20 µl of purified MBP–AtKdos protein in 1 ml was incubated at 37°C for 100 min, and then applied onto a column of Dowex 1X8 (Cl^−^ form, 8 × 20 mm) pre-equilibrated with 20 mM Tris–HCl, pH 8.0. After washing with 20 mM Tris–HCl, pH 8.0, the column was eluted with a linear gradient of 0–0.5 M NaCl in 20 mM Tris–HCl, pH 8.0. Fractions containing KDO8P eluting after Ara5P and PEP were identified by thiobarbituric acid assay (York *et al*., 1985), combined, and desalted on a column of Bio-Gel P-2 (16 × 800 mm, Bio-Rad) equilibrated with 20 mM NH_4_HCO_3_. Fractions containing KDO8P were lyophilized, dissolved in 20 mM Tris–HCl, pH 8.0, and stored at −20°C.

### Subcellular localization of GPP proteins

Constructs for the expression of GPP2::GFP or GPP1::GFP fusion proteins under the control of the CaMV35S promoter were first generated, and their promoters were then replaced with the Arabidopsis *UBQ10* promoter as follows. The coding sequence of *GPP2* was amplified by PCR from an Arabidopsis cDNA library using primer 5 and primer 6. The amplified product was cloned into pENTR/D-TOPO (Invitrogen) and transferred to the pGWB5 vector (Nakagawa *et al*., 2007) using the Gateway technology. The resulting plasmid carrying the construct to express GPP2::GFP under the control of CaMV35S promoter was designated as pGWB5–AtGPP2. A DNA fragment containing the *UBQ10* promoter and a part of *GPP2* cDNA was prepared by a fusion PCR. A DNA fragment containing the upstream sequence of *UBQ10* was PCR-amplified from genomic DNA using *Hin*dIII-linked primer 7 and primer 8 (Grefen *et al*., 2010). A *GPP2* cDNA fragment was PCR-amplified from an Arabidopsis cDNA library with primer 9 and primer 10. The two PCR products were mixed and subjected to PCR with primer 7 and primer 10 to yield the *UBQ*10 promoter fused to *GPP2* partial cDNA. The PCR product was cleaved at the *Hin*dIII site upstream of the *UBQ10* promoter and the *Spe*I site in the *GPP2* coding sequence and then used to replace the *Hin*dIII/*Spe*I region of pGWB5–AtGPP2.

The coding sequence of GPP1 was amplified by PCR from the cDNA library using primer 11 and primer 12, cloned into pENTR/D-TOPO, and transferred to pGWB5 to yield the plasmid pGWB5–AtGPP1 for the CaMV35S promoter-driven GPP1::GFP expression. The *UBQ10* promoter was PCR-amplified from genomic DNA using primer 7 and primer 13. A DNA fragment containing the *GPP1* cDNA and a part of GFP cDNA was PCR-amplified from pGWB5–AtGPP1 using primer 14 and primer 15. The two PCR products were mixed and subjected to a PCR with primer 7 and primer 15 to yield a fusion of the *UBQ10* promoter and the partial GPP1::GFP sequence. The PCR product was cleaved at the *Hin*dIII site upstream of the *UBQ10* promoter and the *Kpn*I site in the *GPP1* coding sequence and then used to replace the *Hin*dIII/*Kpn*I region of pGWB5–AtGPP1.

Agrobacterium-mediated transient expression in Arabidopsis seedlings was performed essentially as described (Li *et al*., 2009; Grefen *et al*., 2010). Arabidopsis seeds were surface-sterilized with 70% ethanol, vernalized, and germinated in 3 ml quarter-strength MS liquid medium (pH 5.8) containing 0.5% sucrose in 6-well culture plates. The day before starting co-cultivation, colonies of *Agrobacterium tumefaciens* GV3101(pMp90) carrying the binary vectors were picked from agar plates and inoculated in 2 ml Luria-Bertani medium containing 20 µg ml^−1^ rifampicin, 50 µg ml^−1^ kanamycin and 50 µg ml^−1^ hygromycin. The next day, 1 ml of saturated culture was diluted with 10 ml of fresh medium containing antibiotics and cultured for 6–8 h. The bacterial cells were harvested by centrifugation at 6,000×*g* for 5 min, washed once with 10 ml of 10 mM MgCl_2_ and 150 µM acetosyringone, and resuspended in a quarter-strength MS liquid medium (pH 7.0). The suspension was adjusted to the density of 1.00–1.25 McFarland turbidity standard with the quarter-strength MS medium (pH 7.0) supplemented with 1% sucrose, 150 µM acetosyringone, and 0.003% (v/v) Silwet L-77 to be used as a co-cultivation medium.

Three to six 4-day-old seedlings were soaked in 3 ml of the co-cultivation medium on a 6-well plate. After incubation in the dark for 40−48 h, seedlings were surface-sterilized with 0.1% sodium hypochlorite for 10 min, washed three times with sterile distilled water, and then incubated overnight in quarter-strength MS medium containing 0.5% sucrose and 500 µg ml^−1^ cefotaxime. Seedlings were stained with MitoTracker Red CMXRos (Cambrex Bio Science Walkersville) and examined for GFP or MitoTracker fluorescence using a confocal laser scanning microscope (FluoView FV500, Olympus).

### Analysis of T-DNA insertion mutant

Seeds of T-DNA insertion lines in the *GPP2* locus (SALK_017815, SALK_062406, and SALK_106665) were obtained from ABRC, and the homozygous T-DNA lines were screened from their progeny by PCR. Primers to identify the insertion lines were designed using the SALK T-DNA verification primer design program (http://signal.salk.edu/tdnaprimers.2.html).

The abundance of *GPP2* transcript in WT and the T-DNA insertion lines was compared using semi-quantitative reverse transcription-PCR. Total RNA was extracted from young leaves using RNeasy Plant Mini kit (Qiagen) according to the manufacturer’s instructions. First-strand cDNA was synthesized using d(T)_18_ primer and ReverTra Ace reverse transcriptase (TOYOBO) and used as the template for PCR with primers AtGPP2-cDNA-Fwd and AtGPP2-cDNA-Rev. *Actin2* (*ACT2*) was amplified with primers ACT2-Fwd and ACT2-Rev as an internal control to calibrate the amounts of cDNA.

Seeds were surface-sterilized and sown on half-strength MS medium containing 1% (w/v) sucrose and 0.8% (w/v) gellan gum (bioWORLD), pH 5.8, unless otherwise stated. After vernalization at 4°C for 2 days, the plates were incubated in a vertical orientation in the growth chamber. Images of seedlings were taken with a scanner, and the primary root lengths were measured using ImageJ software (Schneider *et al*., 2012). Liquid culture was carried out in half-strength MS medium containing 1% (w/v) sucrose and vitamins. Flasks containing 8–10 surface-sterilized seeds and 25 ml of medium were shaken on a rotary shaker (NR-2, Titec) under a 16 h/8 h light/dark cycle at 22°C.

The cell wall samples were prepared as alcohol-insoluble residues from liquid-cultured seedlings. Plant tissues (approximately 1–3 g fresh weight) were ground into a fine powder in liquid nitrogen using a mortar and pestle. The powdered tissue was suspended in approximately five volumes of 80% (v/v) ethanol and centrifuged at 5,800 × g for 15 min at 4°C. The pellet was resuspended in 10 ml of 80% ethanol, centrifuged again, and the washing step was repeated a total of three times. The resulting insoluble residue was further washed once with 10 ml of ethanol and once with acetone. The final pellet was air-dried in a fume hood and used for analysis of KDO and B contents.

### Reverse transcription-quantitative PCR analysis

Total RNA was extracted from seedlings using the Total RNA Extraction Kit Mini (Plant) (RBC Bioscience) according to the manufacturer’s instructions. First-strand cDNA was synthesized from the DNase-treated RNA using d(T)_18_ primer and ReverTra Ace reverse transcriptase. Quantitative PCR was performed with KOD SYBR qPCR Mix (TOYOBO) using Thermal Cycler Dice Real Time System (TP-850, Takara Bio). Expression levels of target genes were normalized against those of *ACT2*. The gene-specific primers used for the analysis are listed in Supplementary Table S1.

### Structural analysis

Multiple sequence alignments of GPP2 with its plant homologs and bacterial KDO8Pases were performed using MAFFT version 7.526 (Katoh and Standley, 2013).

The predicted three-dimensional structure of Arabidopsis GPP2 (AF-Q8VZP1) was obtained from the AlphaFold Protein Structure Database (https://alphafold.ebi.ac.uk/). Structural superposition of the GPP2 core domain and the bacterial KDO8Pase from *Moraxella catarrhalis* (PDB ID: 4UMF) was performed using UCSF ChimeraX (version 1.11.1) (Meng *et al*., 2023).

Molecular docking of KDO8P into the putative substrate-binding pocket of GPP2 was carried out using AutoDock Vina (version 1.2.5) (Trott and Olson, 2010; Eberhardt *et al*., 2021). Among the docking poses, those positioning the phosphate group of KDO8P in proximity to the catalytic Asp residue (Asp20) were selected based on mechanistic considerations. From these, a pose exhibiting a plausible in-line arrangement for nucleophilic attack (Allen and Dunaway-Mariano, 2004) was chosen for further refinement. The selected pose was refined in UCSF ChimeraX by adjusting the ligand orientation to obtain a catalytically plausible configuration, while maintaining reasonable hydrogen-bonding geometry. The refined model was subjected to local energy minimization using AutoDock Vina and evaluated for consistency with the catalytic mechanism of the HAD superfamily.

### Assay methods

Boron was quantified by the chromotropic acid method (Matoh *et al*., 1997). Cell wall KDO contents were estimated by measuring the amount of side chain C of RG-II [rhamnosyl-(1→5)-KDO; Rha-KDO]. An aliquot of cell wall material (1 mg) was suspended in 0.1 M sulfuric acid and heated at 100°C for 30 min to selectively release Rha-KDO from RG-II. In parallel, known amounts of RG-II were subjected to the same hydrolysis procedure to generate a calibration curve. Aliquots of the centrifuged supernatant were mixed with equal volumes of labeling reagent [14 mM 1,2-diamino-4,5-methylenedioxybenzene (DMB; Sigma-Aldrich), 2.8 M acetic acid, 1.4 M 2-mercaptoethanol, and 36 mM sodium hydrosulfite (Nacalai Tesque)] and incubated at 50°C for 2.5 h. The DMB-derivatized samples were analyzed by HPLC with fluorescence detection. Separation was performed on a reversed-phase Unison UK-C18 column (100 × 4.6 mm, Imtakt). The DMB-derivatized Rha-KDO was detected using RF-10AXL fluorescence detector (Shimadzu) with excitation and emission wavelengths set at 373 and 485 nm, respectively. The flow rate was 0.7 ml min^−1^, and the injection volume was 20 µl. Elution was carried out with a linear gradient of acetonitrile:water from 5:95 (v/v) to 15:85 (v/v) over 40 min.

The degree of RG-II dimerization was estimated as reported (Matsunaga and Ishii, 2006) using a column YMC-Pack Diol-120 (800 × 8.0 mm, YMC Co.) equilibrated with 0.2 M NaCl in 50 mM sodium acetate and a refractive index detector RI-8020 (TOSOH).

### Statistical analysis

Statistical significance was determined using Student’s t-test. All calculations were performed using R software (version 4.5.0; R Core Team, 2025).

### Use of AI-assisted writing

Large language models, including ChatGPT (OpenAI) and Gemini (Google), were used for language editing and readability improvement. The models were not used for data generation, analysis, or interpretation. After using these tools, we reviewed and edited the content as needed and take full responsibility for the content of the publication.

## Author contributions

T.H., Y.W., and M.K. performed the research and analyzed the data; M.K. and T.M. designed the research and supervised the study; M.K. acquired funding; T.H. and M.K. wrote the manuscript; all authors reviewed and approved the final version of the manuscript.

## Acknowledgments

This work was supported by JSPS KAKENHI (Grant Numbers 23580090, 26450078, 17K07696, 20K05767, and 24K01655 to MK) and by Grants-in-Aid for Scientific Research on Innovative Areas (Grant Numbers 23119509 and 15H01232 to MK) from MEXT, Japan. We thank Dr. Tsuyoshi Nakagawa (Shimane University) for providing pGWB vectors, Dr. Tetsuro Okuno and Dr. Masanori Kaido (Kyoto University) for the use of the confocal laser scanning microscope, ABRC and the Salk Institute Genomic Analysis Laboratory for providing the sequence-indexed Arabidopsis T-DNA insertion mutants.

## Data availability statement

Data that support the findings of this study are available within the article and its Supporting Information files. Raw data and materials are available upon request.

## Funding statement

This work was supported by JSPS KAKENHI (Grant Numbers 23580090, 26450078, 17K07696, 20K05767, and 24K01655 to MK) and by Grants-in-Aid for Scientific Research on Innovative Areas (Grant Numbers 23119509 and 15H01232 to MK) from MEXT, Japan.

## Conflict of interest

The authors declare no conflict of interest.

## Ethics approval statement

Not applicable

## Patient consent statement

Not applicable

## Permission to reproduce material from other sources

Not applicable

## Supplementary files

**Supplementary Figure S1.**
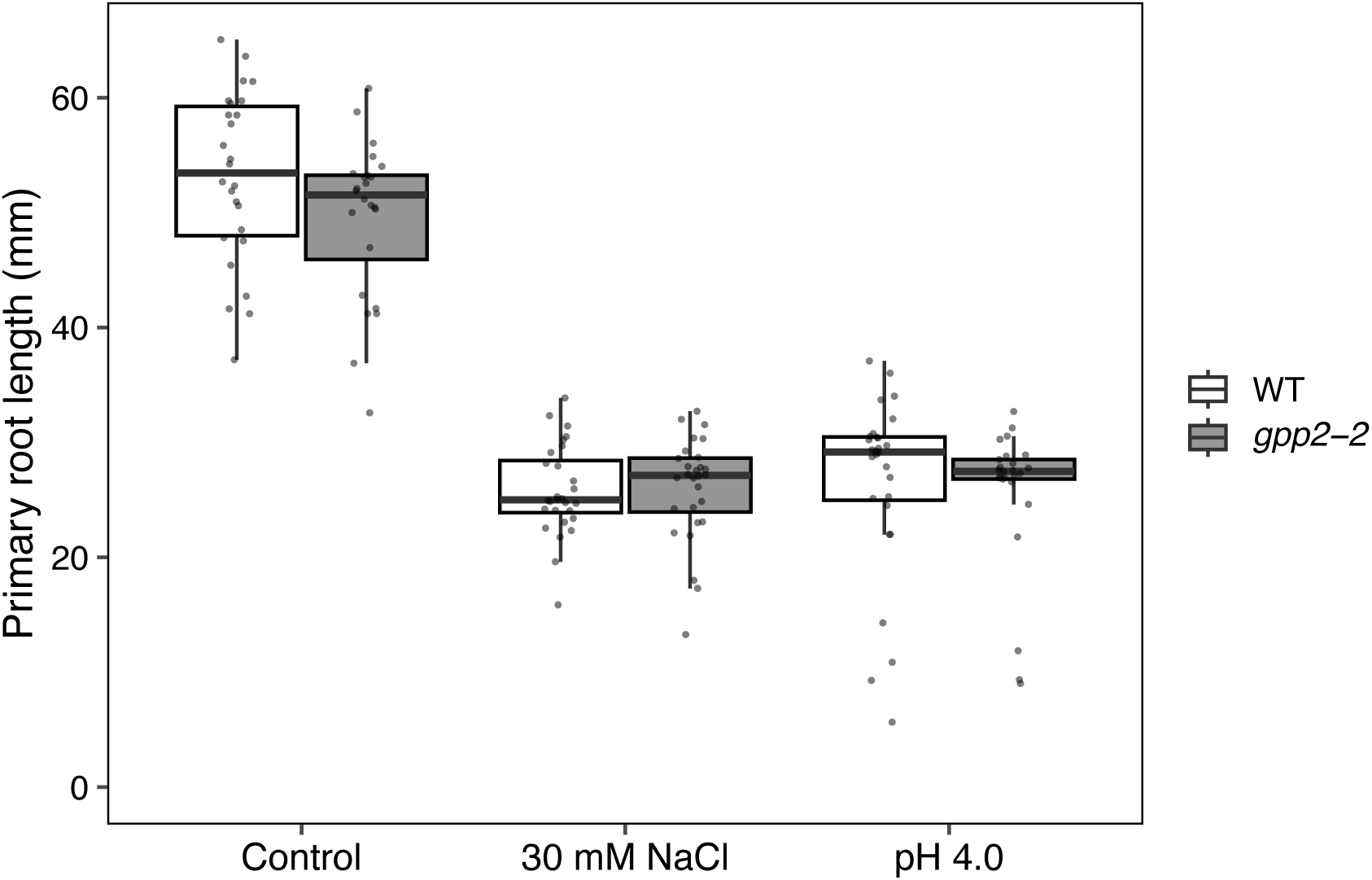
Growth of *gpp2* mutant plants under salt and low pH stresses. Primary root length of wild-type (WT) and *gpp2-2* seedlings grown for 7 days on half-strength MS media (control), or media containing 30 mM NaCl or adjusted to pH 4.0. Individual data points are shown on the boxplots. No significant differences were observed between WT and the mutant under salt or low pH stress conditions (*p* > 0.05; Student’s *t*-test; n = 23–27).

**Supplementary Figure S2.**
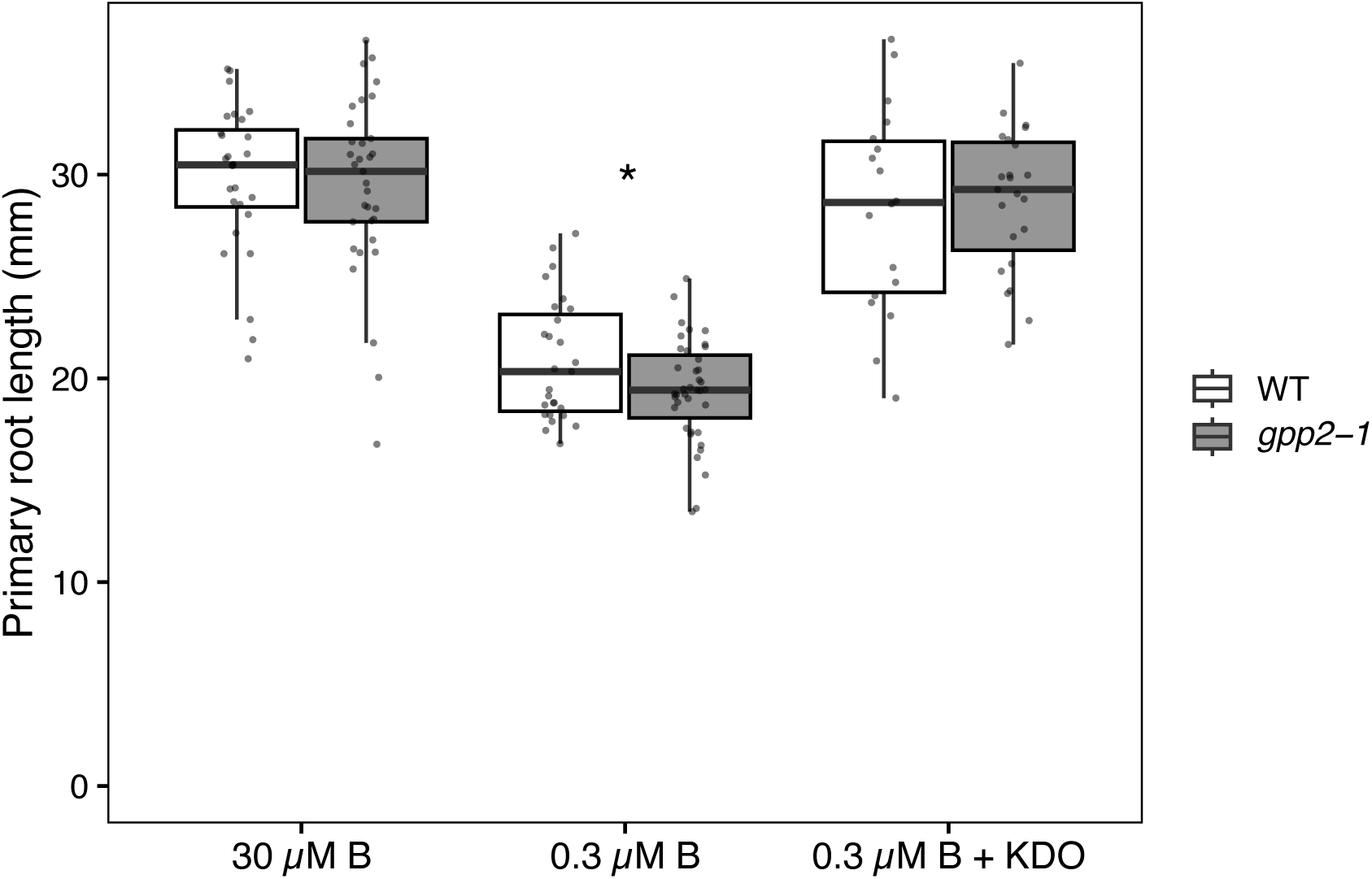
Effect of exogenous KDO supply on the root growth of *gpp2* mutant plants. Primary root length of WT and *gpp2-1* seedlings grown for 7 days on half-strength MS media containing 30 µM B (normal), 0.3 µM B (low-B), or 0.3 µM B supplemented with KDO (low-B + KDO). Individual data points are shown on the boxplots. The asterisk indicates a significant difference between WT and the mutant (*p* < 0.05; Student’s *t*-test; n = 18–39).

**Supplementary Table S1.**
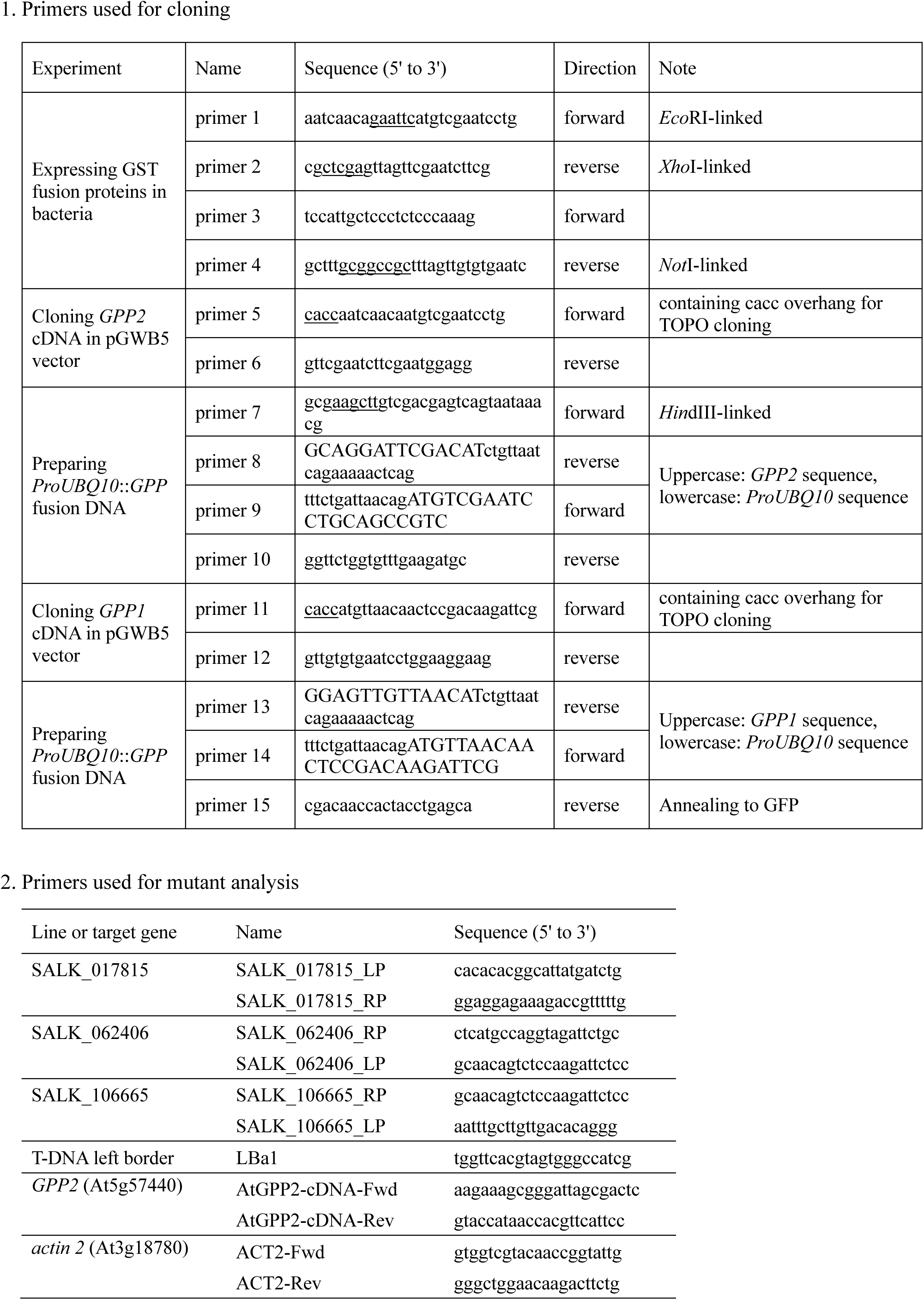
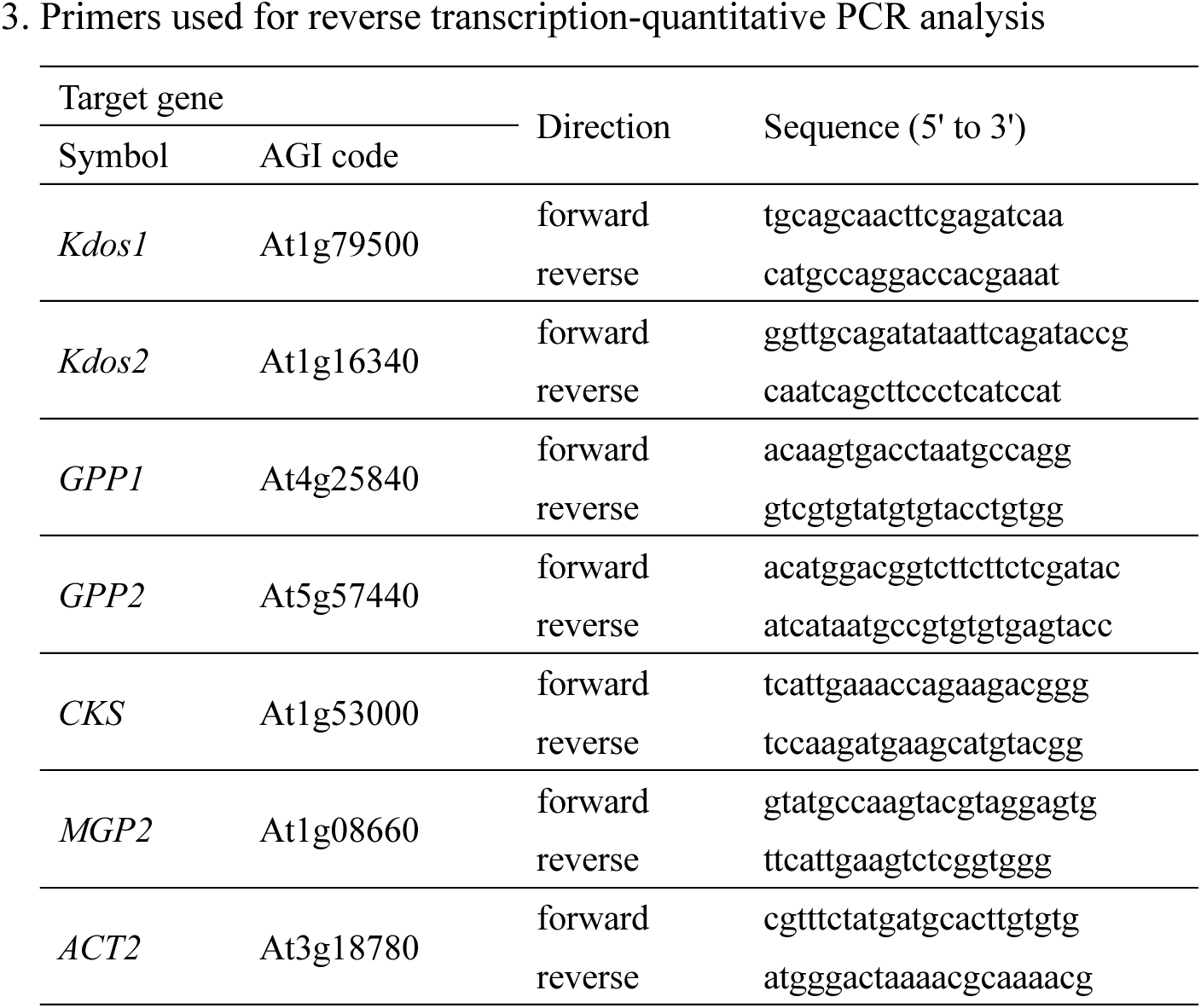
Primers used in this study.

## Notes

### Competing Interest Statement

The authors have declared no competing interest.

